# Recombination and low-diversity confound homoplasy-based methods to detect the effect of SARS-CoV-2 mutations on viral transmissibility

**DOI:** 10.1101/2021.01.29.428535

**Authors:** Elena E. Giorgi, Tanmoy Bhattacharya, Will Fischer, Hyejin Yoon, Werner Abfalterer, Bette Korber

## Abstract

The SARS-CoV-2 variant carrying the Spike protein mutation G614 was first detected in late January 2020 and within a few months became the dominant form globally. In the months that followed, many studies, both in vitro and in animal models, showed that variants carrying this mutation were more infectious and more readily transmitted than the ancestral Wuhan form. Here we investigate why a recently published study by van Dorp et al. failed to detect such higher transmissibility of the G614 variant using homoplasy-based methods. We show that both low diversity and recombination confound the methods utilized by van Dorp et al. and significantly decrease their sensitivity. Furthermore, though they claim no evidence of recombination in their dataset, we and several other studies identify a subset of the sequences as recombinants, possibly enough to affect their statistic adversely.

## Introduction

Here we provide a counterpoint to the recent paper by Van Dorp et al.^1^ entitled, *“No evidence for increased transmissibility from recurrent mutations in SARS-CoV-2,”* where the authors report no evidence for increased transmissibility of any SARS-CoV-2 variant using a phylogenetic strategy based on recurrent mutations. At the time of their publication (11/25/2020), however, there was a substantial body of evidence that at least one mutation in the Spike (S) D614G mediates phenotype that is substantially more infectious, and more readily transmitted, than the ancestral Wuhan form. This variant and its derived lineages have essentially replaced the ancestral form that initiated the global spread from China and now dominate the global pandemic (Fig. S1); multiple studies have experimentally demonstrated increased infectivity and transmissibility of viruses carrying the mutation S D614G (this mutation, which defines the “G” clade, is usually accompanied by 3 additional co-transmitted mutations outside of Spike). Therefore, one should ask not, “Does the S D614G mutation confer a more infective phenotype?” but rather, “How could the method employed by van Dorp et al. fail to detect it?” We believe the method fails due to two interrelated effects: first, limited diversity (Fig. 1) and undetected recombination events (Fig. 2) undermine the reliability of the SARS-CoV-2 phylogenetic reconstruction; second, recurrent mutations simply do not provide a sensitive test for detecting selection in this evolutionary context.

**Fig. 1.**
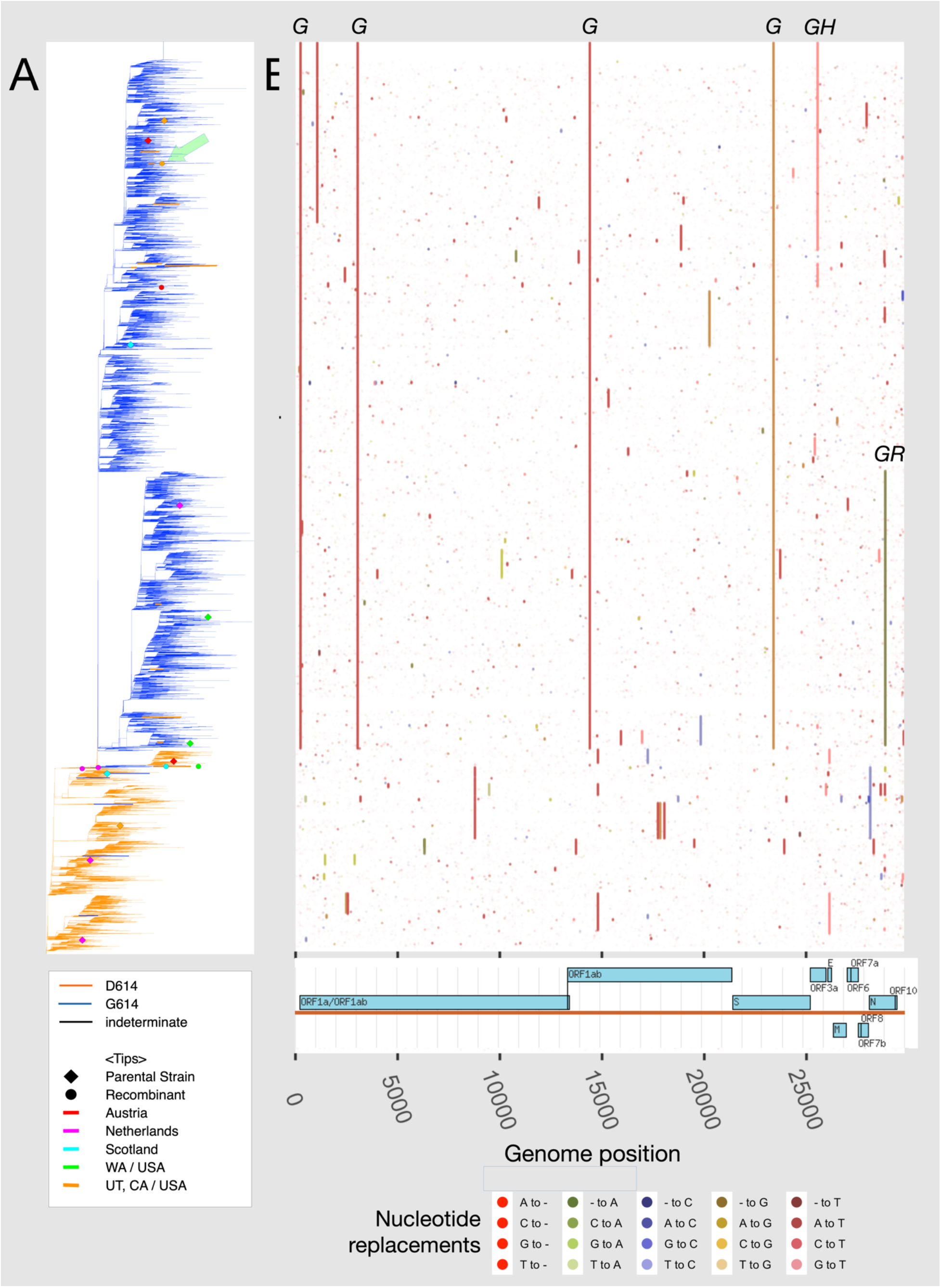
Mutations in SARS-CoV-2 genomes comprising the van Dorp *et al.* dataset, arranged according to the van Dorp *et al.* phylogenetic tree. (**A)** A revisualization of the phylogenetic tree from van Dorp *et al.* (data from their Fig 1). Here, we color branches by the amino-acid present at position S 614: orange for the original outbreak amino-acid (asparagine, D), blue for the replacement (glycine, G). Black denotes sequences where the amino acid at 614 was undetermined. Putative recombinant sequences and inferred parental strains are marked with colored circles and diamonds, respectively, and correspond to those defined in Figure 2. The G clade emerges at the point where the G614 amino acid indicated by the blue branches begins to dominate; there are some recombination events in the region of the tree near the origin of the G clade that likely arise similarly to the situation illustrated in detail in Fig. S2. The D614 mutation is very rarely found in the context of the G clade; where present, its origin might be back-mutation, but also possibly undetected recombination, or sequencing artifacts. In one such case, we found evidence of a recombinant origin for the UT/CA sequence shown in Fig. 2C (green arrow). **(B)** A matrix showing mutation type and location, spanning the full SARS-CoV-2 genome from left to right on the x-axis, for the 46,773 full sequence alignment in the van Dorp *et al.* data set, compared to an ancestral-identical isolate, GISAID accession EPI_ISL_418509. The sequences in the matrix are tree-ordered, corresponding to the phylogeny provided by van Dorp on the left. The 4 linked mutations that are associated with the G clade are seen in this matrix as the long vertical stripes associated with the clade that generally carries S D614G (C241T, C3037T, C14408T, and A23403G). The alignment and tree here are based on data provided by van Dorp from their publication, and incorporate their masking of 68 internal sites (i.e. all bases in masked columns are converted to N, so variation is ignored) following De Maio et al.^20^. Mutations defining the “G”, “GH”, and “GR” clades are marked at the plot margin.

**Fig. 2.**
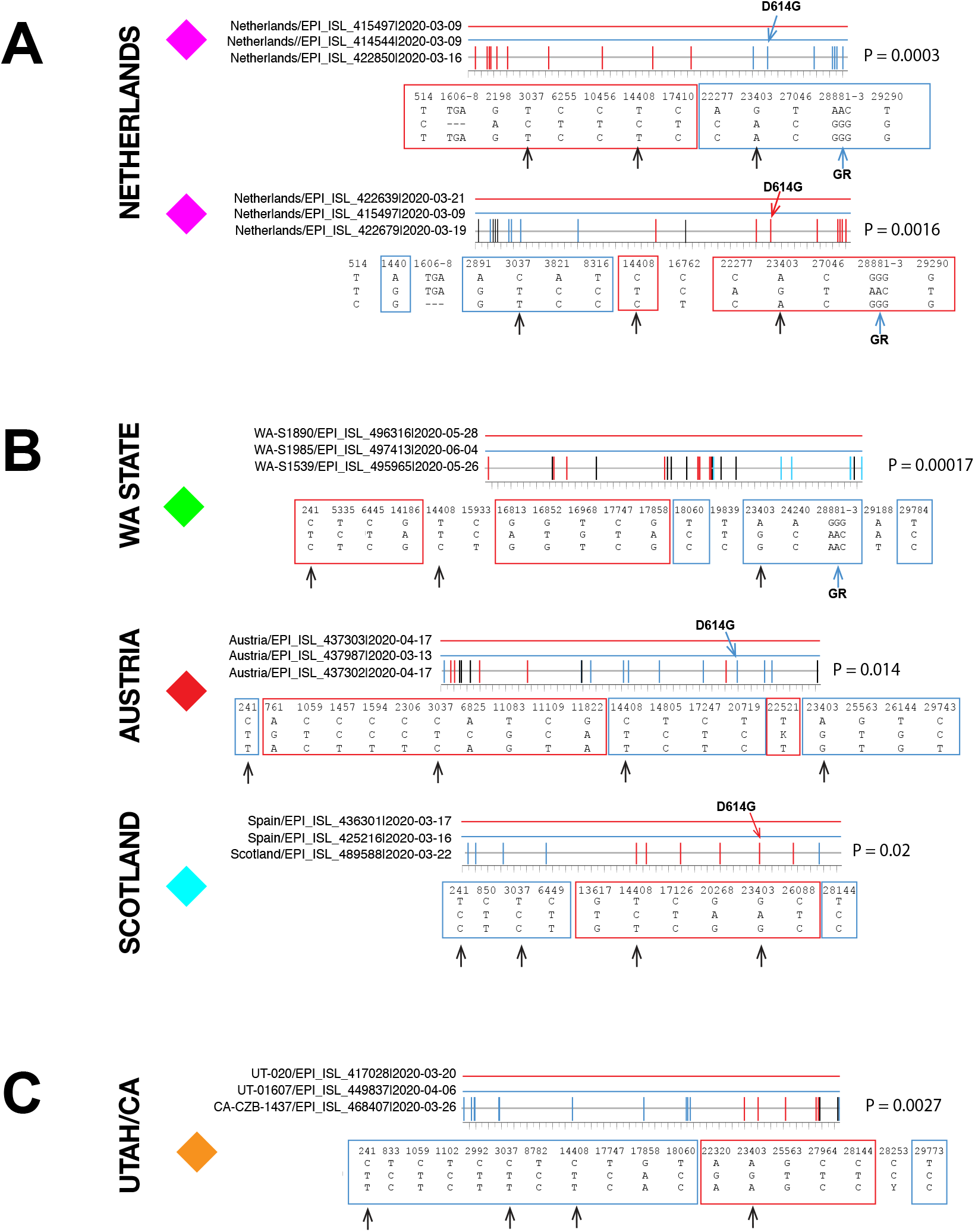
Examples of recombinants found in within the van Dorp alignment from WA state, the Netherlands, Austria, and Scotland. Each graph represents a recombination event detected by the software RAPR^24^. Parental strains are shown in red (top line) and blue (second line), and the child is shown below with color-coded tic-marks representing mutations matching either parental strain. Nucleotides in black do not match either parent. Boxes below show nucleotides at each position of diversity across the triplet, with red or light blue boxes highlighting the parental strain they match. Black arrows show the four D614G G-clade defining positions: C241T (the UTR was not included in the Netherlands alignment), C3037T, C14408T and A23403G (D614G in the spike gene), and, when relevant, a fourth arrow indicates the triplet of consecutive mutations that characterizes the GR clade at (28881-3). Each of these recombination sets can be found in the tree in Figure 1 marked by diamonds (parents) and circles (recombinants) in the same color shown here on the left of each set; this is the same set of sequences used by Van Drop where they find no recombination. **(A)** Two recombinants from the Netherlands. We looked for recombination specifically in the Netherlands as a region of interest because we had prior evidence of rare disruptions of major clades within the datasets. All triplet combinations of sequences from these regions were compared, and the P-values are shown from the Wald–Wolfowitz runs test, which were significant after an FDR correction for multiple tests^24^. A more detailed example of one of these local recombination sequence histories, and how recombination disrupts the phylogenetic reconstruction of the entire set, is shown in Figure S2. (Of note, VanInsberghe et al.^22^ did not find evidence for these recombination events applying their methods. This was at least in part because they use only limited sites that were characteristic of the major clades for the identification of recombinants; these sites have limited overlap with the set of sites that define the particular recombination events we show here. In contrast, in our analysis, we utilize information from all sites and our approach is based on evaluating all mutational patterns in all triplets among locally cocirculating sequences^22^.) **(B)** Three more examples of recombinants found in different countries, namely, Washington (USA), Austria, and Scotland (with parental strains from Spain). These sets were evaluated because of their position in van Dorp’s tree, essentially between the ancestral form and the emergence of the G clade, as seen in Figure 1 (how this situation can arise is illustrated in Fig. S2). In these examples, the 4-base haplotype of the G clade is disrupted, and additional variant positions found among geographically local sequences provide statistical support for recombination. We used the actual sequences for a given isolate to explore recombination, not the masked sets used by van Dorp as shown in Figure 1, to include more information and increase power. **(C)** As a final example, we show a recombinant from California, originally reported by both VanInsberghe et al.^22^ and Varabyou et al. ^23^, which was also confirmed by our RAPR method24. We display the most probable assignment of the parent-child relationship in each recombination triplet, given the background set under study, and parents themselves may be part of a recombinant lineage; additional data may be able to better inform the directional assignments within a triplet. A key point here is that positive evidence of recombination can be found in among the sequences van Dorp studied where they found no evidence of recombination, but different methods that use different strategies will find different examples. By including rare variants in our analysis, we can find examples that utilize this information that will be missed by other methods. Other methods, however, can find recombinants in the global set that we miss, because we are constrained to look in more local geographic regions, as exploring all of GISAID to too computationally intensive and suffers from severe multiple testing limitations.

## Methods

Phylogenetic reconstructions were based on input data provided by van Dorp et al. The additional sequence sets use to detect recombinants were obtained from the Global Initiative for Sharing All Influenza Data (GISAID) database (Elbe and Buckland-Merrett, 2017; Shu and McCauley, 2017) and tracked using the tools provided in the LANL COVID-19 Viral Genome Analysis Pipeline (available at https://cov.lanl.gov/content/index) as previously described^2^.

To identify the candidate recombination parent and child sequences shown in Fig. 2 and S2, we first enumerated all hypotheses about recombination triplets (i.e., two putative parents and a child). For each triplet, we ignored all columns in the alignment that had any gaps, ambiguity codons, or multiple states, and focused on the columns remaining that had a mutation. We filtered out the hypotheses where the child had more unique mutations than similarities to either parent; for the remaining we calculated the p-value against the clustering being explainable as arising from random mutations using the Wald Wolfowitz runs test statistic (Wald and Wolfowitz, 1940). We then computed the Benjamini-Hochberg critical value (Benjamini and Hochberg, 1995) for the false discovery rate to be less than 10%. In this analysis, we ignored the statistical nonindependence of the tests due to the underlying phylogenetic dependence of the tests. A web interface for to run the RAPR code for is available at www.hiv.lanl.gov. The code to enable multiple test corrections of large data sets is available through GitHub.

The selection analysis presented in the supplemental figure is based on the methods we used in our previous publication^2^.

## Results

### Evidence for increased transmissibility of SARS-CoV-2 variants carrying Spike (S) D614G

The rapidity and *highly repetitive* nature of the transition in prevalence from the ancestral form of SARS-CoV-2 to the S D614G form, observed at every geographic level in every region of the world once the forms were co-circulating, was, of itself, indicative of a selective advantage^2^; the overall consistency of this trajectory can be seen in Supplemental Fig. S1. Multiple lines of experimental evidence indicate that the viruses with the S D614G mutation are more infectious *and* more transmissible than the ancestral form. First, patients infected with viruses carrying this mutation required lower RT-PCR cycle thresholds (Ct) for detection, indicating higher viral loads in the upper respiratory tract^3–6^. Second, the G614 Spikes were significantly (3-to 6-fold) more infectious than the ancestral D614 form, in multiple, independent sets of distinct pseudotype viral assays^2,7,8^. Finally, viruses with the S D614G mutation transmit faster and outcompete the ancestral virus *in vivo* in animal models^9^, and the authentic virus carrying the S D614G is more infectious in cultured human large airway epithelial cells, even when the ancestral virus is seeded in 10-fold excess^9^. In addition, the S D614G mutation increases sensitivity to neutralization by convalescent sera, vaccine sera, and some monoclonal antibodies^10^. Structural^7,10,11^ and molecular dynamics studies^12^ revealed that the S D614G mutation confers a preference for a more open “up” conformation, increasing the exposure of the ACE2 receptor binding site, as well as key epitopes for neutralizing antibodies^10^, offering a mechanistic explanation of the greater infectivity and neutralizing sensitivity of this variant.

### Recombination

Coronavirus infections are frequent and widespread across different animal reservoirs, where they co-exist in the same hosts and often recombine^16^. The 2015 human MERS-CoV virus, for example, was a recombinant lineage from different MERS-CoV species circulating camels. Furthermore, there have been reports of several occurrences where it recombined with a co-circulating human CoV 229E lineage^19^, reflecting the fact that coronaviruses are prone to recombination within and across different coronavirus species. The recombinatory origin of SARS-CoV-2 has been well documented^15,19^. More importantly, despite its short circulating time and still relatively low diversity, more and more studies are documenting occurrences of recombination within the SARS-CoV-2 rapidly emerging new clades^25,26^. Van Dorp *et al.* state, “*we find no signature of recombination in SARS-CoV-2*”. Lack of a significant regression of linkage disequilibrium with distance between loci argues for a rarity of recombination events, not for their absence. Recombination is clearly an important aspect of coronavirus evolution ^13–16^, which has been shown to play a role in the origins of SARS-CoV-2^15,17,18^ and SARS-CoV-1^19^. More directly relevant, several studies have found occurrences of recombination among SARS-CoV-2 variants^20–23^. Two of them find evidence of recombination in SARS-CoV-2 based on major clades and their defining mutations^22,23^. We used a method that enables us to also utilize rarer mutations; as this approach is computationally intensive, we focused on variants co-circulating in geographically local sets (Fig. 2)^24^. We find strong support for recombination among some sequences used in van Dorp, highlighted in Fig. 1 and detailed in Fig. 2. Recombination can cause problems in phylogenetic reconstruction. Varabyou *et al.* identified 225 plausible recombinants among 87,695 GISAID SARS-CoV-2 genomic sequences, and illustrated their potential for distorting the phylogenetic tree^23^; many of these would be expected to overlap with the data set sampled from GISAID used by van Dorp et al. for their analysis. In Supplement Fig. S2, we provide an example illustrating one of several ways a recombinant SARS-CoV-2 sequence can be improperly placed in a phylogenetic tree. Recombination frequently causes atypically long branch lengths and may give rise to temporal anomalies; de Maio *et al.* identified 30 putative recombinant SARS-CoV-2 sequences associated with these characteristics, and removed them from subsequent analyses^20^. Given that the methods use different strategies for identifying recombinants, they would naturally identify distinct sets of recombinants within the dataset; here our intent is just to identify sequences within this data set that are likely recombinants. Given the slow accumulation of point mutations in SARS-CoV-2, the potential even for rare recombination events to impact its evolution should not be underestimated. Most problematically for the van Dorp *et al.* analysis, spurious placements of recombinant sequences can disrupt otherwise homogeneous clades, corrupting the sister-clade variant distributions that their statistic requires. One particularly clear example of this is a recombinant sampled in California that was identified using both clade-based approaches^22,23^ as well as using our strategy (Fig. 2C)^24^, and that gives rise to a D614 variant embedded within the G clade (Fig 1A). While recombinant sequences clearly arise naturally in coronaviruses, they may also occur as *in vitro* artifacts resulting from strand switching during PCR amplification^20,25^, or as sequence assembly miscalls of minor variants^23^. Regardless of their origin (in nature, test tube, or software), recombinants violate methodological assumptions, and complicate phylogenetic inference and interpretation of trees; thus, it is important to be aware that recombination can be found among SARS CoV-2 sequences.

### Insensitivity of homoplasy-based strategies for identifying positive selection in settings of limited diversity

Van Dorp and colleagues attempted to assess evidence for positive selection in the SARS-CoV-2 genome quantifying the relative number of descendants in sister clades after a homoplasy event to identify increased transmissions. In situations where the accumulation of point mutations is slow, as in SARS-CoV-2 (Fig. 1B), the number of times a mutation appears in an unlinked context (*i.e., de novo*) is also small, seriously limiting statistical power. Van Dorp *et al.* mention this lack of power, and their confidence intervals for the D614G mutation allow for the possibility of substantial positive selection (van Dorp’s Fig. 3). While they failed to find statistical support for positive selection for *any* mutation recurring in the global GISAID dataset, including D614G in Spike, the absence of such evidence is not evidence against positive selection. The major reconstructed sister clades they define are occasionally disrupted in terms of the amino acid at 614 (Fig. 1). The 4 bases of the G clade, which includes the D614 mutation, are linked and this linkage is very rarely disrupted (Fig 1B, also detailed in Korber et al.^2^, supplementary Fig. S5); in the few cases when it is disrupted, it may be due to *de novo* mutations, but also perhaps to undetected recombination, mixed bases in consensus calls, or poor phylogenetic resolution in this low diversity setting. In their attempt to avoid overcounting when applying their phylogenetic index, van Dorp *et al.* do not analyze such disrupted clades: therefore, the rare disruptions profoundly impact their test statistic, and they only consider very limited sampling near a few tips of the phylogenetic tree. Thus, for example, the very large difference in the size of the two clades separated by the D614G mutation near the base of the tree does not contribute to their statistic.

## Discussion

New clades and mutations continue to emerge that are potentially under positive selection including some that are spreading rapidly in the United Kingdom (UK) (B.1.1.7) ^26^, South Africa (501Y.V2) ^27,28^ and Brazil (484K.V2)^29^ and such variants of interest need to be experimentally evaluated to explore their impact on infectivity and immunological sensitivity. Early detection of variants of interest to test for possible phenotypic transitions in the virus through sequence analysis is the first critical step to ensure that COVID-19 countermeasures continue to be effective. In low-diversity settings of recent epidemics caused by slowly mutating pathogens, homoplasy-based methods, or other methods based on analysis of phylogeny alone, are not expected to have the power to detect selective advantage. In contrast, use of additional geographic or population data, such as evidence of consistent replacement of a clade by another in multiple independent sub-epidemics, is more likely to uncover ongoing selection. This will require maintaining a steady effort on global surveillance of diversity, and reversing the recent decline in sequence sampling and submission to GISAID from many parts of world (Fig. S1); hopefully vaccine roll-out will stimulate more extensive sampling in these under-sampled regions. Methods to experimental assay variants of interest advanced greatly throughout 2020, and now can be readily deployed, allowing sequence-based epidemiological evidence for the potential importance of a site to be rapidly linked to experimental validation. Indeed such experimental validation was undertaken for the D614G mutation and provided strong evidence for increased infectivity^2^ and transmissibility^30^. Further, it is important to understand that recombinant sequences are evident among the global SARS COV-2 data sets, and recombination can impact interpretation of the evolutionary history of SARS-CoV-2 variants ^23^.

## Acknowledgments

We thank van Dorp and colleagues for their open sharing of sequence alignments and phylogenetic trees, enabling a direct comparison to their work. We were supported by the Laboratory Directed Research and Development program of Los Alamos National Laboratory under project number ECR 20200554ECR.

## Competing interests

The authors declare no competing interests.

## Author contributions

B.K. conceived the response and drafted the manuscript; B.K., E.E.G., W.F. and T.B. analyzed the data, prepared the figures, and edited the manuscript; H.Y., W.F, and W.A. contributed to the analyses.

## Supplemental Figures

**Fig. S1.**
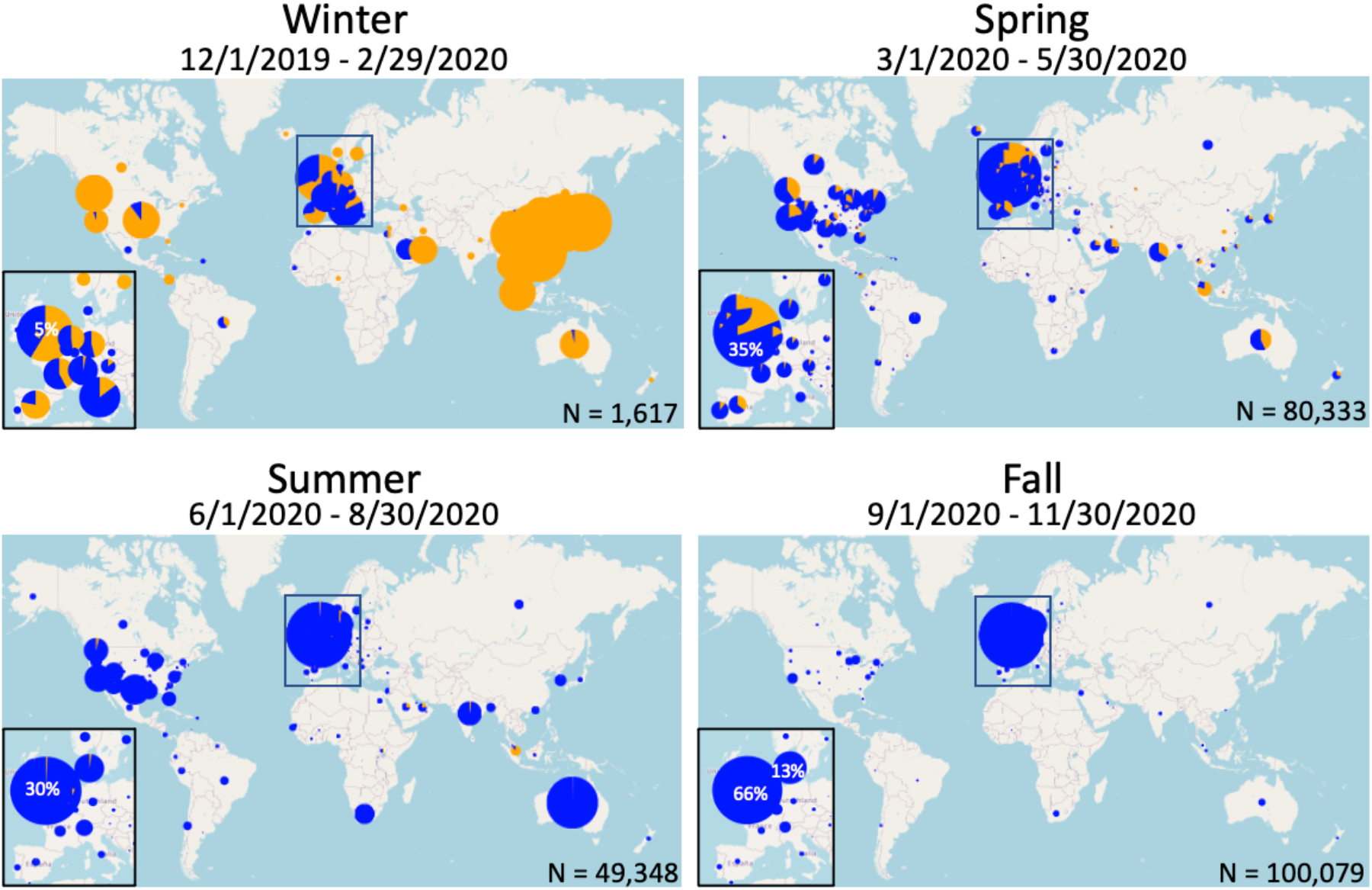
Maps of GISAID data over 3 month windows showing the global transition from the more ancestral form of the virus with S D614 (orange) to viruses carrying the S D614G form. The size of the circle indicates the relative sampling size within each map, the number of sequences (N) for each time window is noted at the bottom right. A blow up of Europe is shown in the bottom left, as samples from England are relatively so enriched they obscure sampling from other regions in Europe. This figure is updated daily from GISAID at cov.lanl.gov, and this map is based on GISAID data included in the “regional complete” alignment of Spike, extracted and aligned as described in Sup. Fig. 1 of Korber *et al.*^2^ The number of sequences available in each time window is indicted at the bottom right. The fraction of the total sequences that were sampled in the United Kingdom (UK) are indicated in white in the circle representing the UK. In the fall panel the fraction of sequences coming from Denmark are also indicated; Denmark and the UK combined accounted for 79% of the global sample. They include available GISAID sequences the span intact coding regions for: Spike, ORF3a, nsp3, and RdRp. The Spike COMPLETE alignment used here includes all intact sequences that encompass the Spike gene NC_045512 bases, 21,563-25,384. The sequences are passed through a quality control filter for inclusion that primarily filters out incomplete sequences, and was based on a Jan. 12, 2021 GISAID data set^2^.

**Fig. S2.**
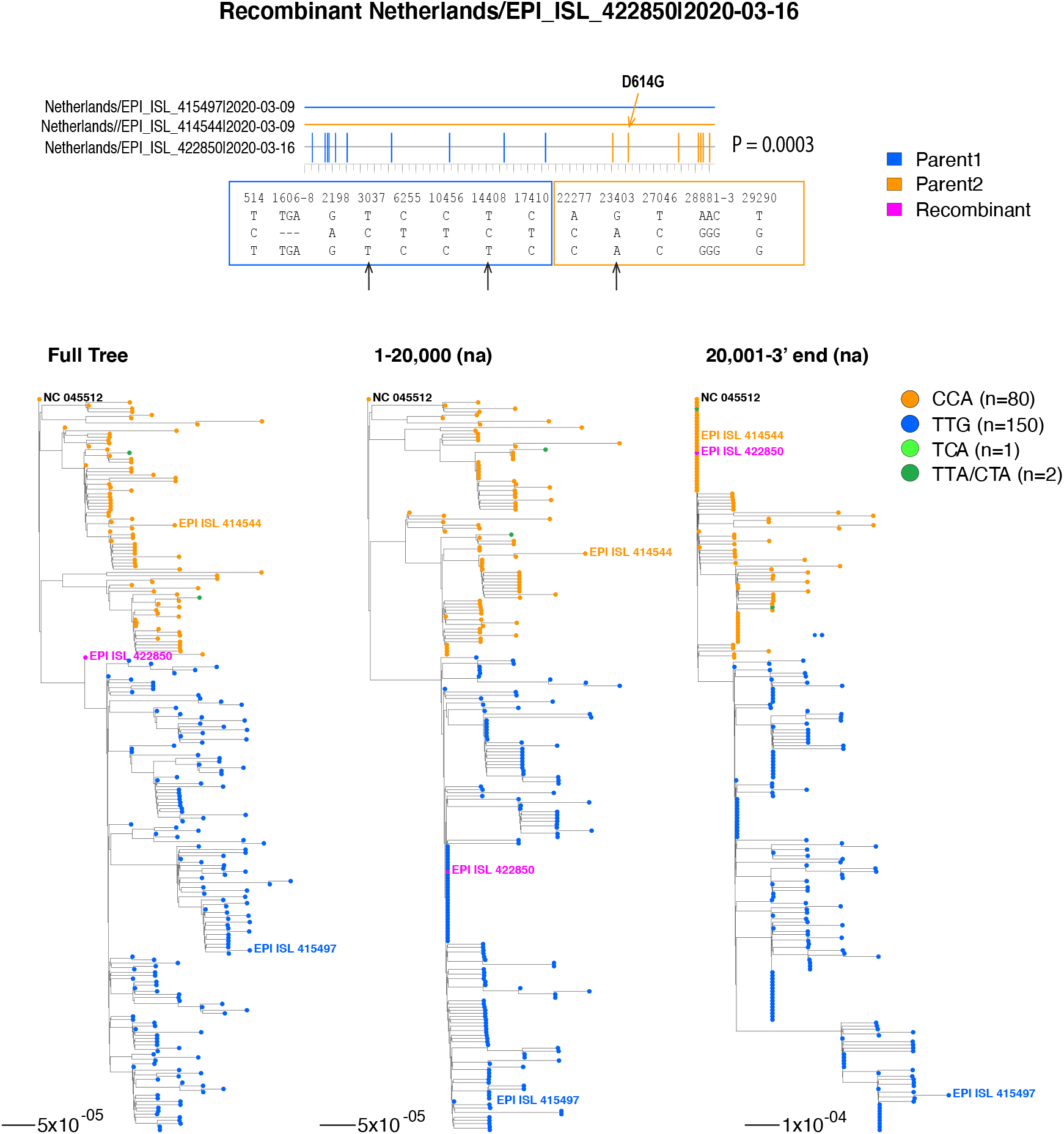
An example showing of how recombinants can affect tree phylogenies. One of the two Netherland recombinants from Fig. 2 is shown again here to illustrate how recombination can disrupt the topology of phylogenetic trees. Top: highlighter plot of recombinant triplet, with parental strains shown in blue (G614 variant) and orange (D614 variant). Mutations found in each sequence are shown below, within color-coded boxes: blue for the 5-prime half, matching the G614 parental variant, and orange for the 3-prime half, matching the D614 parental variant. Bottom left: full tree of 132 unique SARS-CoV-2 sequences from the Netherlands (retrieved from GISAID in April 2020) from which the recombinant triplet shown at the top was found, rooted on the Wuhan reference sequence. The recombinant triplet sequences are shown in bold, with the recombinant shown in magenta. Middle and right panels: separate phylogenetic trees of the 5-prime and 3-prime halves of the genome, split in the region where the recombination breakpoint likely occurred. In the middle tree the recombinant falls in the G614 clade, where it matches the G614 parent, whereas in the tree on the right it falls in the D614 clade, matching the second parent.

## Notes

https://www.nature.com/articles/s41467-020-19818-2

### Competing Interest Statement

The authors have declared no competing interest.

## References

1 van Dorp, L. et al. No evidence for increased transmissibility from recurrent mutations in SARS-CoV-2. Nat Commun 11, 5986, doi:10.1038/s41467-020-19818-2 (2020).

2 Korber, B. et al. Tracking Changes in SARS-CoV-2 Spike: Evidence that D614G Increases Infectivity of the COVID-19 Virus. Cell 182, 812–827, doi:10.1016/j.cell.2020.06.043 (2020).

3 Lorenzo-Redondo, R. et al. A Unique Clade of SARS-CoV-2 Viruses is Associated with Lower Viral Loads in Patient Upper Airways. medRxiv, 2020.2005.2019.20107144, doi:10.1101/2020.05.19.20107144 (2020).

4 Mueller, N. F. et al. Viral genomes reveal patterns of the SARS-CoV-2 outbreak in Washington State. medRxiv, doi:10.1101/2020.09.30.20204230 (2020).

5 Volz, E. et al. Evaluating the effects of SARS-CoV-2 Spike mutation D614G on transmissibility and pathogenicity. Cell, doi:https://doi.org/10.1016/j.cell.2020.11.020 (2020).

6 Wagner, C. et al. Comparing viral load and clinical outcomes in Washington State across D614G mutation in spike protein of SARS-CoV-2. (2020).

7 Yurkovetskiy, L. et al. Structural and Functional Analysis of the D614G SARS-CoV-2 Spike Protein Variant. Cell 183, 739–751.e738, doi:10.1016/j.cell.2020.09.032 (2020).

8 Zhang, L. et al. The D614G mutation in the SARS-CoV-2 spike protein reduces S1 shedding and increases infectivity. bioRxiv, 2020.2006.2012.148726, doi:10.1101/2020.06.12.148726 (2020).

9 Hou, Y. J. et al. SARS-CoV-2 D614G variant exhibits efficient replication ex vivo and transmission in vivo. Science, eabe8499, doi:10.1126/science.abe8499 (2020).

10 Weissman, D. et al. D614G Spike Mutation Increases SARS CoV-2 Susceptibility to Neutralization. medRxiv, 2020.2007.2022.20159905, doi:10.1101/2020.07.22.20159905 (2020).

11 Gobeil, S. M. C. et al. D614G mutation alters SARS-CoV-2 spike conformational dynamics and protease cleavage susceptibility at the S1/S2 junction. bioRxiv, 2020.2010.2011.335299, doi:10.1101/2020.10.11.335299 (2020).

12 Mansbach, R. A. et al. The SARS-CoV-2 Spike Variant D614G Favors an Open Conformational State. bioRxiv, doi:10.1101/2020.07.26.219741 (2020).

13 Graham, R. L. & Baric, R. S. Recombination, Reservoirs, and the Modular Spike: Mechanisms of Coronavirus Cross-Species Transmission. Journal of Virology 84, 3134–3146, doi:10.1128/jvi.01394-09 (2010).

14 Lai, M. M. & Cavanagh, D. The molecular biology of coronaviruses. Adv Virus Res 48, 1–100, doi:10.1016/s0065-3527(08)60286-9 (1997).

15 Li, X. et al. Emergence of SARS-CoV-2 through recombination and strong purifying selection. Sci Adv 6, doi:10.1126/sciadv.abb9153 (2020).

16 Su, S. et al. Epidemiology, Genetic Recombination, and Pathogenesis of Coronaviruses. Trends Microbiol 24, 490–502, doi:10.1016/j.tim.2016.03.003 (2016).

17 Flores-Alanis, A., Sandner-Miranda, L., Delgado, G., Cravioto, A. & Morales-Espinosa, R. The receptor binding domain of SARS-CoV-2 spike protein is the result of an ancestral recombination between the bat-CoV RaTG13 and the pangolin-CoV MP789. BMC Res Notes 13, 398, doi:10.1186/s13104-020-05242-8 (2020).

18 Lam, T. T.-Y. et al. Identifying SARS-CoV-2-related coronaviruses in Malayan pangolins. Nature 583, 282–285, doi:10.1038/s41586-020-2169-0 (2020).

19 Hon, C.-C. et al. Evidence of the Recombinant Origin of a Bat Severe Acute Respiratory Syndrome (SARS)-Like Coronavirus and Its Implications on the Direct Ancestor of SARS Coronavirus. Journal of Virology 82, 1819–1826, doi:10.1128/jvi.01926-07 (2008).

20 De Maio, N. et al. Issues with SARS-CoV-2 sequencing data (2020).

21 Korber, B. et al. Spike mutation pipeline reveals the emergence of a more transmissible form of SARS-CoV-2. bioRxiv, 2020.2004.2029.069054, doi:10.1101/2020.04.29.069054 (2020).

22 VanInsberghe, D., Neish, A., Lowen, A. C. & Koelle, K. Identification of SARS-CoV-2 recombinant genomes. bioRxiv, 2020.2008.2005.238386, doi:10.1101/2020.08.05.238386 (2020).

23 Varabyou, A., Pockrandt, C., Salzberg, S. L. & Pertea, M. Rapid detection of inter-clade recombination in SARS-CoV-2 with Bolotie. bioRxiv, doi:10.1101/2020.09.21.300913 (2020).

24 Song, H. et al. Tracking HIV-1 recombination to resolve its contribution to HIV-1 evolution in natural infection. Nat Commun 9, 1928, doi:10.1038/s41467-018-04217-5 (2018).

25 Salazar-Gonzalez, J. F. et al. Deciphering human immunodeficiency virus type 1 transmission and early envelope diversification by single-genome amplification and sequencing. J Virol 82, 3952–3970, doi:10.1128/JVI.02660-07 (2008).

26 Davies, N. G. et al. Estimated transmissibility and severity of novel SARS-CoV-2 Variant of Concern 202012/01 in England. medRxiv, 2020.2012.2024.20248822, doi:10.1101/2020.12.24.20248822 (2020).

27 Tegally, H. et al. Emergence and rapid spread of a new severe acute respiratory syndrome-related coronavirus 2 (SARS-CoV-2) lineage with multiple spike mutations in South Africa. medRxiv, 2020.2012.2021.20248640, doi:10.1101/2020.12.21.20248640 (2020).

28 Wibmer, C. K. et al. SARS-CoV-2 501Y.V2 escapes neutralization by South African COVID-19 donor plasma. bioRxiv, 2021.2001.2018.427166, doi:10.1101/2021.01.18.427166 (2021).

29 Naveca, F. et al. Phylogenetic relationship of SARS-CoV-2 sequences from Amazonas with emerging Brazilian variants harboring mutations E484K and N501Y in the Spike protein Virological.org (2021).

30 Hou, Y. J. et al. SARS-CoV-2 D614G Variant Exhibits Enhanced Replication <em>ex vivo</em> and Earlier Transmission <em>in vivo</em>. bioRxiv, 2020.2009.2028.317685, doi:10.1101/2020.09.28.317685 (2020).

